# SARS-CoV-2 JN.1 reveals attenuated pathogenicity and airborne transmission

**DOI:** 10.1101/2024.11.07.622580

**Authors:** Ruixue Liu, Qiushi Jin, Wenqi Wang, Cheng Zhang, Hang Zhang, Bing Li, Fang Yan, Xianzhu Xia, Jianmin Li, Xuefeng Wang, Yuwei Gao

## Abstract

JN.1 is a subvariant of SARS-CoV-2 Omicron BA.2.86 lineage that was predominant worldwide in early 2024, of which the in vivo characteristics are largely unknown. Our results demonstrated that the replication of JN.1 was more efficient than that of the parental BA.2 in Vero cells, which demonstrated low dependence on TMPRSS2. Compared to Omicron variants BA.2 and XBB EG.5.1, JN.1 replicated less efficiently in hACE2 mouse lungs of which the intranasal infection was not lethal to hACE2 mice and led to weaker immune dysregulation. On a more sensitive, aged hACE2 hamster model, JN.1 led to a lower mortality rate and no weight loss, corresponding well with the low preference in lower airways. Lower amounts of viruses in nasal washes and exhaled aerosols were detected in JN.1 infected wildtype hamsters than EG.5.1, and consistently, JN.1 also exhibited largely reduced airborne transmission. Moreover, the poor transmission was also clearly demonstrated even by using hamsters expressing hACE2 receptors in the whole airway. Thus both pathogenicity and airborne transmission of JN.1 were demonstrated to be largely attenuated.

**Importance:** Currently, SARS-CoV-2 JN.1 and its subvariants have fully replaced the previous dominant XBB lineage around the world. Although the strong immune evasion of JN.1 has been distinctly revealed, its in vivo pathogenicity and airborne transmission remained unclear. By using multiple Omicron-sensitive rodent models, our findings demonstrated that the pathogenicity of JN.1 was largely attenuated. The weak airborne transmission of JN.1 in wildtype and hACE2 hamsters was consistent with the reported relative lower transmissibility in human, and the using of airway-expressing hACE2 hamsters ulteriorly eliminates the potential bias in viral transmission studies induced by receptor divergence between animal models and human. These findings uncover the in vivo virological characteristics of SARS-CoV-2 novel lineage, providing insights for communicable disease control.

## Introduction

From late 2023, Omicron BA.2.86 lineage of severe acute respiratory syndrome coronavirus 2 (SARS-CoV-2) with its subvariants began to outcompete the previously dominant XBB lineages worldwide. Compared to BA.2 lineage, BA.2.86 harbors more than 30 mutations in the spike protein, which exhibits antigenicity shift and is identified as a new serotype compared to XBB (1, 2). The BA.2.86 spike protein also reveals higher hACE2 binding affinity than that of BA.2 (1, 3, 4). Both characteristics are regarded to lay molecular foundations to the prevalence of BA.2.86 lineage. Further evolution of the BA.2.86 lineage resulted in its descendant strain, JN.1 (BA.2.86.1.1), which contains a spike-protein substitution L455S. JN.1 rapidly spread globally and was identified as the currently circulating variant of interest (VOI) by World Health Organization in December, 2003. Compared to BA.2.86, JN.1 exhibits robust immune evasion and has become one of the most prevalent variants by now (5, 6).

Syrian hamsters are highly suspectable to SARS-CoV-2 infection and show similar pulmonary pathological features to those of coronavirus disease 2019 (COVID-19) patients after infection, which have been commonly used for SARS-CoV-2 pathogenicity and transmission studies (7–9). Early Omicron VOIs (such as BA.1) demonstrates largely attenuated pathogenicity in a hamster model which is in accord with the clinical reports (10–12). To better reveal the in vivo virological characteristics of Omicron VOIs, more suspectable, multiple hACE2 transgenic rodent models have been commonly applied (11, 13). Recently, BA.2.86 demonstrated lower pathogenicity than both EG.5.1 and BA.2 (14). However, to date, pathogenicity and transmissibility of JN.1 were largely unclear. Here, using H11-K18-hACE2 mice and hamsters which are highly sensitive to Omicron infection (15, 16), we report the pathogenicity and airborne transmission JN.1

## Results

### Prevalence and in vitro replication of JN.1

The prevalence of JN.1 started in the end 2023, which soon outcompeted XBB VOIs, EG.5.1 and HK.3 in North American, Asian and European countries (Fig. 1A) and is keeping circulating globally until now (in August, 2024). We compared the in vitro replication of clinical isolates of JN.1 and BA.2 in Vero E6, Vero E6^TMPRSS2+^, and HeLa^hACE2+^ cells (Fig. 1B). Viral titers of JN.1 were consistently higher than BA.2 in Vero E6 cells from 24 to 96 hours post infection (HPI). Both JN.1 and BA.2 replicated efficiently in Vero E6^TMPRSS2+^ cells, with the highest titers up to about 10^6^ PFU/ml. Interestingly, by comparing the highest titers on Vero E6 and Vero E6^TMPRSS2+^ cells, it was obvious that the growth of BA.2 was largely enhanced (about 33 folds) when TMPRSS2 were expressed, while only 1.43 folds of enhancement was observed for the growth of JN.1 (Fig. 1C). In HeLa^hACE2+^ cells that do not express TMPRSS2 (15), their replications were similar. Revealed by viral loads, TMPRSS2 overexpression also induced weak enhancement of JN.1 replication. The results suggest that entry of JN.1 is less dependent on TMPRSS2 compared to BA.2.

**Figure 1.**
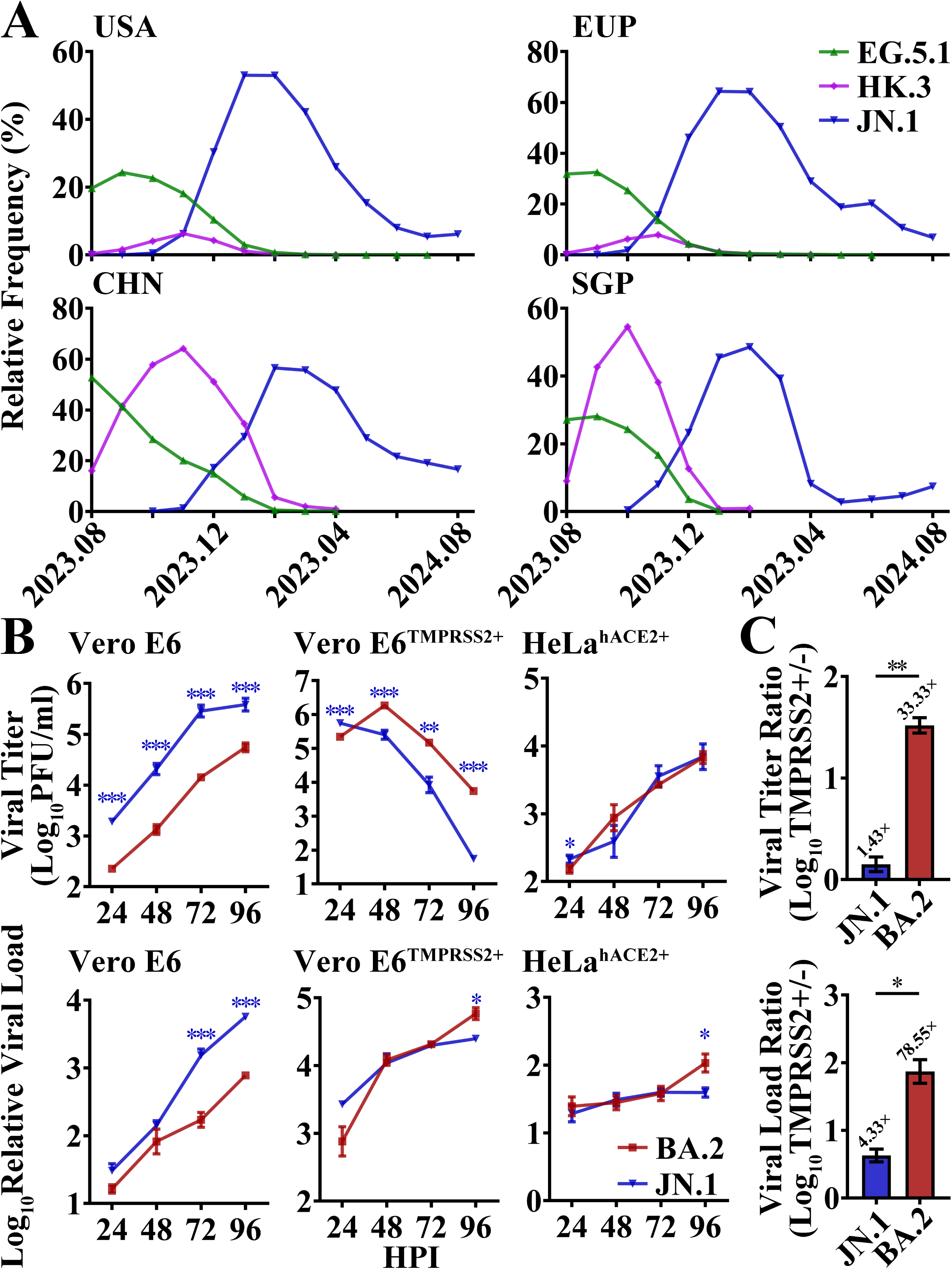
Prevalence and replicative kinetics of JN.1. **A.** Prevalence of EG.5.1 (green), HK.3 (purple) and JN.1 (blue) in China (CHN), the United States of America (USA), Europe (EUP) and Singapore (SGP) for twelve months from August 2023 (2023.08) to August 2024 (2024.08). **B**. Replicative kinetics of BA.2 (dark red), JN.1 (blue) in terms of viral titer (upper) and viral load (lower) in Vero E6, Vero E6^TMPRSS2+^ and HeLa^hACE2+^ cells. **C**. Ratio of replicative efficiencies on Vero E6^TMPRSS2+^ to those on Vero E6 cells. The highest viral titers (upper) or viral loads (lower) of JN.1 and BA.2 in Vero E6^TMPRSS2+^ and Vero E6 cells were used for the analysis. Significance of the differences in replication between BA.2 and JN.1 is indicated above the lines by bule asterisks.

### Pathogenicity in H11-K18-hACE2 mice

We first characterized the pathogenicity of JN.1 using H11-K18-hACE2 mice, in which the infection of an Omicron BA.5.2.48 subvariant led to lethality (15). Intranasally inoculation of a parental Omicron variant, BA.2, led to death of 1/5 mice at 8 days post infection (DPI), and no more death was observed in the following two days (Fig. 2A). The infection of another VOI, EG.5.1, caused a total of 3/5 death by 10 DPI. By sharp contrast, JN.1 infection did not result in lethality in the 10 days observation. Similarly, EG.5.1 infection led to robust body weight loss from 7 to 10 DPI, and the average weight loss reached up to 13% at 9 DPI (Fig. 2B). Nevertheless, insignificant, quiet slight weight loss was revealed in either BA.2 or JN.1 infected mice, although JN.1 mice demonstrated a relative lower weight compared to the mock at 7 DPI. Thus we suggest lower pathogenicity of JN.1 than EG.5.1. Viral loads in oral swabs of JN.1, EG.5.1 and BA.2 groups were comparable at 2, 4, and 6 DPI, respectively, while all of them were approaching undetectable until 8 DPI (Fig. 2C). Revealed by viral titers, JN.1 replicated less efficiently than both EG.5.1 and BA.2 in the lungs of hACE2 mice (Fig. 2D). The viral loads in turbinates of JN.1 mice were also lower than those of EG.5.1 (Fig. S1). The pathology of lungs at 3 DPI were evaluated as previously (16). JN.1 infected lungs showed lower scores than EG.5.1 (Fig. 2E). In details, EG.5.1 infected mice revealed severe thickening of the alveolar wall in a wide range (Fig. 2G). Eosinophilic mucus and epithelial cells necrosis were commonly observed in bronchiole. Focal pulmonary hemorrhage and inflammatory cell infiltration around blood vessels and alveolus were also occasionally observed. In contrast, the infection of JN.1 induced thickening of the alveolar walls and inflammatory cell infiltration to a lesser extent. Gene dysregulation induced by different strains were evaluated by RNA-sequencing. Totally, infection of EG.5.1, BA.2 and JN.1 led to dysregulation of 937, 518 and 281 genes, respectively. Compared to EG.5.1, infection of JN.1 led to less severe upregulation of multiple cytokines including interferon alpha (IFNA1, IFNA2 and INFA9), beta (IFNB1) and lambda (IFNL2 and IFNL3), interleukins (IL6 and IL15) and a granulocyte colony stimulating factor (CSF3) with an antiviral defense gene MX1.

**Figure 2.**
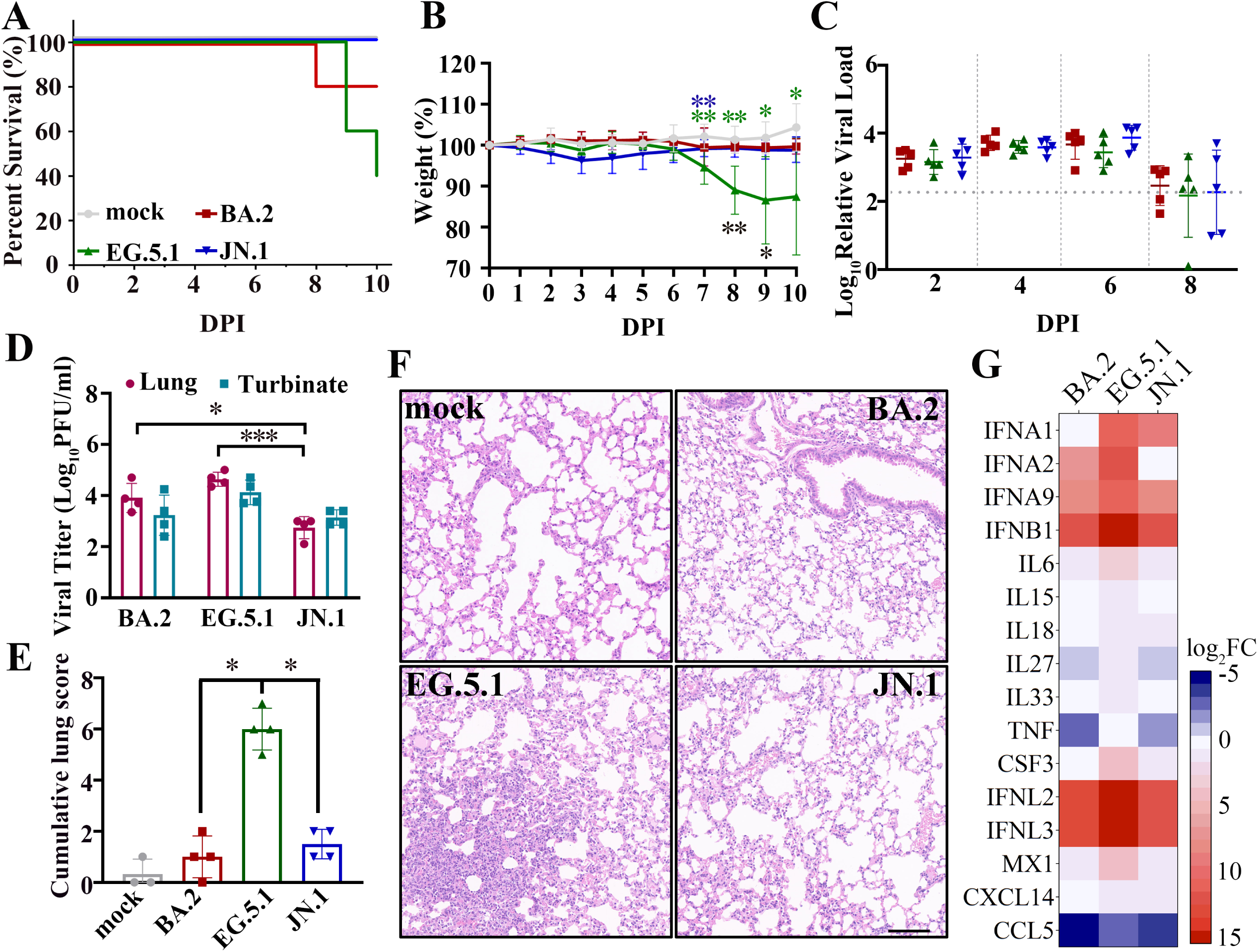
Pathogenicity of JN.1 in H11-K18-hACE2 transgenic mice. K18-hACE2 transgenic mice were intranasally inoculated with BA.2, EG.5.1, or JN.1, respectively. Five mice per group were used to measure the respective parameters (in A, B and C). Four mice per group were euthanized at 3 DPI and used for data collection (in D, E, F and G). The data (in A, B, C and E) of the mock, BA.2, EG.5.1, and JN.1 groups are represented in gray, red, green, and blue, respectively (as shown in A). **A**. Percentage survival. **B.** Percentage body weight changes. Significance of differences between the mock group and each infected group are revealed above the lines using asterisks in colors corresponding to the respective infected group. Significance of differences between JN.1 and EG.5.1 groups are revealed below the lines using black asterisks. **C**. Viral loads in nasal washes. The detection limit is represented by a gray dotted line. **D**. Viral titers in the lungs (dark red) and turbinates (cray). **E**. Cumulative pathological scores of lungs. **F**. H&E staining sections of the lungs. The scale bar represents 100 μm. **G**. Expression profile of infected lungs. Compared to the mock, up- or down-regulation of the genes in infected groups are indicated by negative or positive log_2_fold changes (log_2_FC) in blue and red, respectively.

### Pathogenicity in H11-K18-hACE2 hamsters

In order to clearly characterize the pathogenicity of JN.1, we tended to use H11-K18-hACE2 hamsters which were reported to be Omicron-lethal (15, 16) and thus highly suitable for Omicron pathogenicity studies. Exactly speaking, we here used 52-week-old hamsters, as aged animals were reported to be more sensitive to SARS-CoV-2 infection (17, 18). EG.5.1 infection led to 1/4 death as early as 5 DPI and then 4/4 death by 6 DPI (Fig. 3A), while hamsters infected by JN.1 demonstrated 3/4 lethality by 6 DPI. Body weight loss induced by EG.5.1 infection was revealed as early as 4 DPI, which reached up to 10% at 5 DPI (Fig. 3B). By sharp contrast, no significant weight loss of JN.1 group was observed, while significant differences in body weight changes were clear demonstrated between JN.1 and EG.5.1 as early as 2 DPI. The two groups showed no differences of viral titers in nasal washes at 2 DPI (Fig. 3C). From 2 to 4 DPI, infectious viruses in nasal washes dramatically decreased, and the infectious viruses of 1/4 of EG.5.1 nasal washes and 3/4 JN.1 nasal washes were hardly visible at 4 DPI. The viral loads in nasal washes revealed similar results (Fig. S2A). In homology to the results revealed in hACE2 mice, viral titers and viral loads in the lungs of the JN.1 group were lower than those of EG.5.1 (Figs. 3D and S2B).

**Figure 3.**
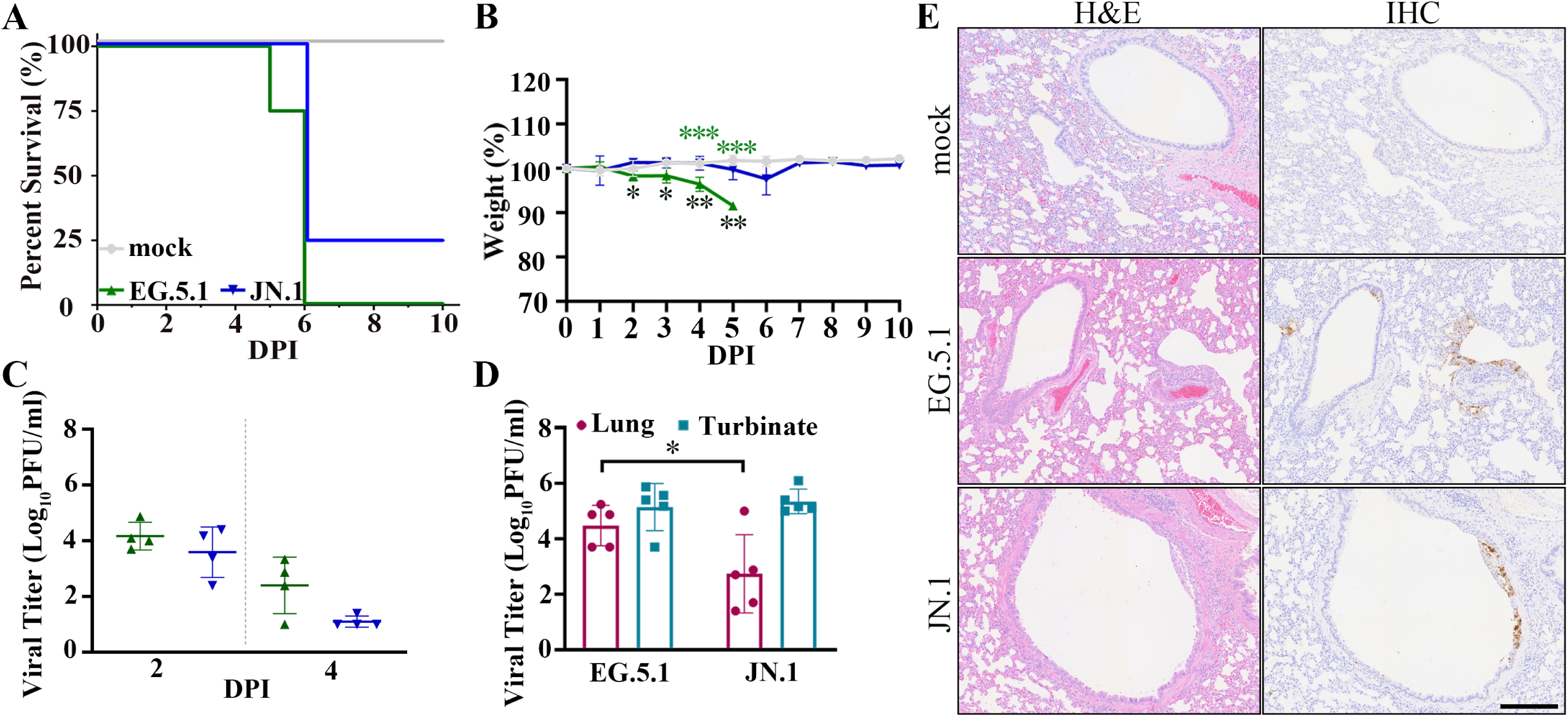
In vivo virological characteristics of JN.1 in K18-hACE2 aged hamsters. H1-K18-hACE2 aged hamsters were intranasally inoculated with EG.5.1 or JN.1, and corresponding results (A-C) of mock-, EG.5.1 and JN.1 infected groups are shown in gray, green and blue, respectively (as shown in A). **A**. Percentage survival. **B**. Percentage body weight changes. Significance of differences between the mock group and each infected group are revealed above the lines using asterisks in colors corresponding to the respective infected group. Significant differences between JN.1 and EG.5.1 groups are revealed below the lines using black asterisks. **C**. Viral titers in nasal washes. **D**. Viral titers in the lungs (dark red) and turbinates (cray). **E**. Histopathology of hamster lungs revealed by H&E staining (left) and IHC (right). Scale bar: 250 μm.

To evaluate the pathological features and determine the infected areas in lungs, we performed H&E staining and anti-nucleocapsid immunohistochemistry (IHC) of the hamster lungs. Similar to the results revealed by hACE2-mouse models, both EG.5.1 and JN.1 infections led to bronchial epithelium necrosis and defluxion and inflammatory cell infiltration around bronchioles and terminal bronchioles (Fig. 3E). However, EG.5.1, other than JN.1 efficiently infected alveolar sacs and alveoli, suggesting that JN.1 has a lower preference to the lower respiratory tract, even when hACE2 receptor is expressed.

### Airborne transmission

The airborne transmission is a key driving force of the epidemicity of SARS-CoV-2 multi-subvariants, which was commonly evaluated using hamster models (19–21). Compared to the poor airborne transmission of the initial Omicron VOIs, such as BA.1 and BA.2, airborne transmissibility of recent VOIs including BA.5.2.48, XBB.1.5 and XBB.1.9 (including EG.5.1 and HK.3 subvariants) were largely enhanced (15, 16, 19, 21, 22). To reveal the comparative airborne transmission of JN.1 and EG.5.1, one donor hamster was firstly nasally inoculated with the indicated virus 24 hours prior to the co-house with one corresponding exposed acceptor. The co-house was maintained for 24 hours to allow airborne transmission. In accord with previous results (16, 21), EG.5.1 was detected in 5/5 turbinates of the acceptors (Fig. 4A), thought infectious viruses were detected only in two lungs. Unexpectedly, although JN.1 replicated efficiently in airways of donors, infectious virus was not detected in any of the acceptors. Considering that the potential low affinity between the spike protein of JN.1 and hamster ACE2 receptor potentially lead to the poor transmission in wildtype (WT) hamsters, the H11-K18-hACE2 hamsters which express hACE2 receptors in turbinate, trachea, and lung (Fig. S3) were applied to characterize the transmission. For EG.5.1, the transmission was detected in 4/5 acceptors (Fig. 4B), while only one acceptor was infected during the JN.1 transmission, of which low amount of infectious virus (just above the detection limit) was detected in the turbinate. Thus our results in WT and hACE2 hamsters suggest the intrinsic attenuated airborne transmission of JN.1.

**Figure 4.**
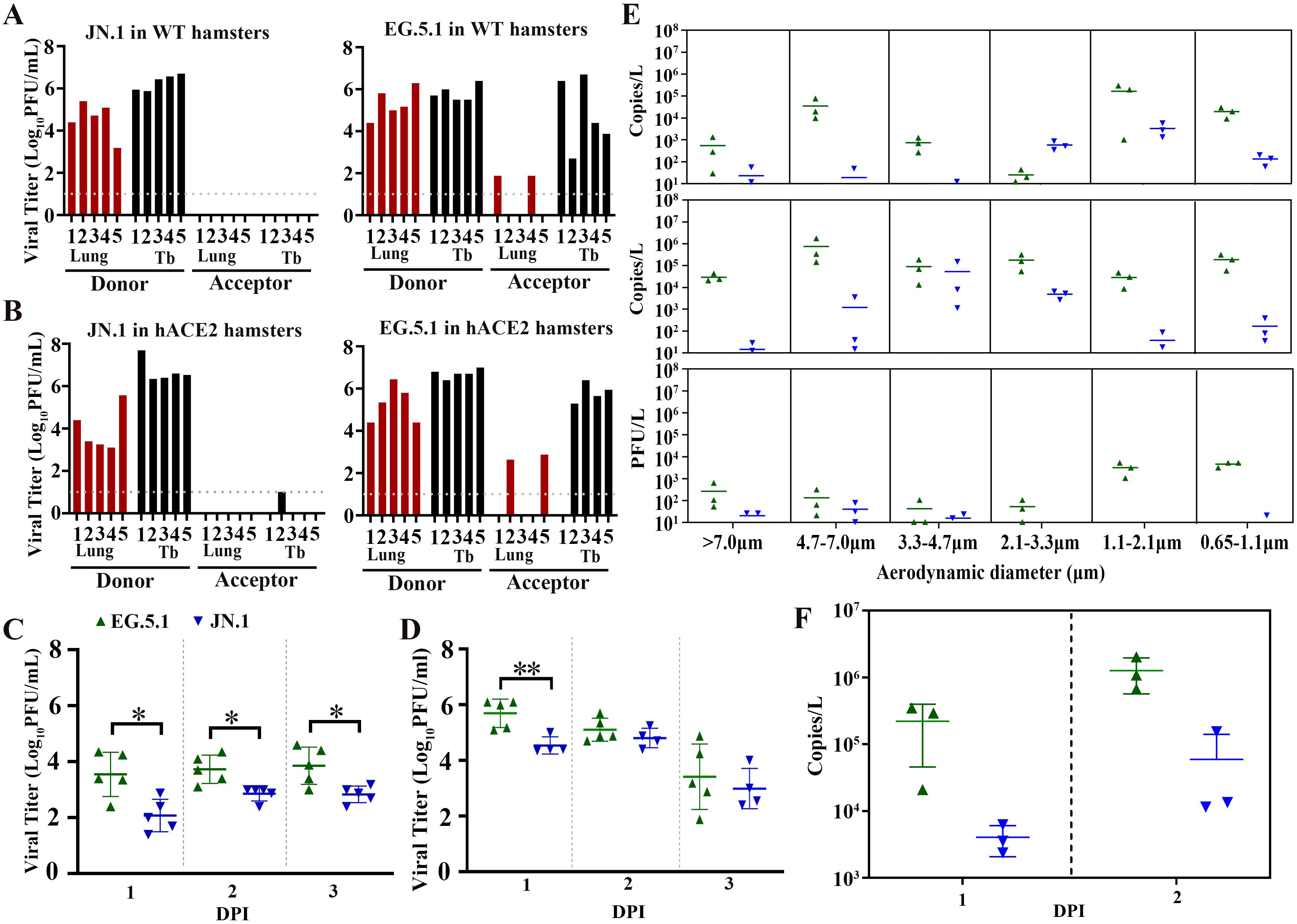
Airborne transmission of JN.1 in hamsters. **A-B**. Airborne transmission of JN.1 and EG.5.1 in WT (A) or H11-K18-hACE2 (hACE, B) hamsters. The viral titers in the lungs and turbinates are indicated in dark red and black, respectively. **C-D.** Time-course viral titers in nasal washes of infected WT (C) and hACE2 (D) hamsters. **E**. Size distribution of viral loads (at 1 DPI in the upper panel, and at 2 DPI in the middle panel) and infectious virus at 2 DPI (lower panel) in exhaled viral particle-loaded aerosols of infected WT hamsters. **F**. Total amounts of viruses in the exhaled aerosols at 1 and 2 DPI.

To investigate the key reason of the attenuated transmission, we further compared the viral titers in nasal washes of JN.1 and EG.5.1 infected WT and hACE2 hamsters. In nasal washes of WT hamsters, viral titers of JN.1 kept at least 40 folds lower than those of EG.5.1 from 1 to 3 DPI (Fig. 4C). When hACE2 were expressed in upper airways of hamsters, viral titers in nasal washes were largely increased (from ∼10^3^ PFU/ml to ∼10^5^ PFU/ml), indicating that the hACE2 hamster is a potential more sensitive, unbias animal model to evaluate SARS-CoV-2 transmission. However, in hACE2 hamsters, much fewer infectious viruses were still revealed in nasal washes of JN.1 groups at 1 DPI (Fig. 4D). Similarly, we also observed lower viral loads in nasal washes of JN.1 infected hamsters (Fig. S4). In exhaled aerosols which were collected totally from 4 hamsters, it is obvious that viral loads in both small particles (with areodynamic diameters of 0.65 to 2.1 μm) and large particles (with diameters over 4.7 μm) of JN.1 aerosols were much lower than those of EG.5.1 at both 1 and 2 DPI (Fig. 4E, upper and middle panels). At 2 DPI, infectious viruses in small particles of EG.5.1 aerosols kept over 1000 PFU/L, which were distinctly higher than those of JN.1 aerosols (Fig. 4E, lower panel). Consistently, the viruses in total aerosols (including small, middle and large particles) of JN.1 were 85 and 42 folds lower than those of EG.5.1 at 1 and 2 DPI, respectively (Fig. 4F). Collectively, lower amounts of exhaled viruses were clearly revealed in JN.1 infected animals, which was in highly positive correlation with the attenuated airborne transmission. In order to reveal the differences of the infected upper airways, the nasal turbinates at 2 DPI were decalcified, and pathological features and infected areas were compared. Infection of EG.5.1 strongly destroyed cilia structure in epithelium and even led to total disappearance of cilia and necrosis of the epithelium on the outer side, while intact cilia was observed from inside to outside in JN.1 infected nasal (Fig. S5). Thus, although both EG.5.1 and JN.1 efficiently infected epithelium in inferior nasal meatus, it was suggested that infection of JN.1 to epithelium in upper airway was much milder than that of EG.5.1.

## Discussion

Previous studies have revealed that pathogenic features in hACE2 transgenic rodent models accords with clinical symptoms in humans, and that initial Omicron VOIs demonstrate relative lower pathogenicity (11, 12). However, pathogenicity of Omicron VOIs do not always keep decreasing. Compared to BA.2, mutations in spike protein of BA.5 were considered to enhance pathogenicity (23). Our previous findings also revealed the enhanced pathogenicity of BA.5.2.48 and XBB.1.9 variants (including previous VOIs, EG.5.1 and HK.3) than BA.2 (15, 16). However, compared to BA.2 and EG.5.1 variants, infection of JN.1 was not lethal in hACE2 mice and even failed to induce weight loss in more-sensitive aged hACE2 hamsters. Moreover, JN.1 induced mild immune dysregulation and revealed a low preference and low efficient replication in lung. All the facts suggest that the pathogenicity of JN.1 is largely attenuated. Clinical reports demonstrated that alveolar damage and subsequently induced systemic hyperinflammation commonly predicted COVID-19 severity and survival (24), which accords well with the pathogenicity in the multiple Omicron-sensitive hACE2 animal models.

JN.1 displayed significantly enhanced immune escape compared to the parental BA.2.86, which was considered to be a key fact to promote its prevalence (5, 6). Besides the immune evasion, the transmissibility also serves as an important driving force to the prevalence. Compared to EG.5.1, the exhaled viruses were much less and the airborne transmission of JN.1 was largely attenuated. Furthermore, by using hamsters overexpressing hACE2 in the whole airway, it was demonstrated that the reduced transmission was not due to the unrecognition of ACE2 receptor in animal models. Our results correspond well with the lower effective reproduction number (Re) of JN.1 in the initial state of prevalence than that of EG.5.1 (5, 25), indicating that JN.1 exhibits potential relative low transmission in human.

Interestingly, our results revealed that both JN.1 and EG.5.1 replicated efficiently in turbinates of WT and hACE2 hamsters, which seem not in accord with the relative low transmission of JN.1 at first glance. However, a previous study on influenza virus indicated that the tropism to nasal epithelium, other than the replication in the whole upper airway, highly affects the viral airborne transmissibility (26). Both EG.5.1 and JN.1 reveal tropism to epithelium in inferior nasal meatus, while severe infection induced by the infection of EG.5.1 other than JN.1 suggest that EG.5.1 replicated better in epithelium of upper airways. Consequently, a weak tropism to nasal epithelium was regarded to partially contribute to the attenuated airborne transmission of JN.1, at least in hamsters. As the latest VOI, JN.1 is still prevailing worldwide currently and keeping mutating swiftly, of which its subvariants LB.1, KP.2.3 and KP.3.1 have demonstrated both increased immune evasion and higher Re number than JN.1 (27). Thus careful surveillance and virological studies of JN.1 with the subvariants provide direct insights for SARS-CoV-2 control and treatment.

We note two limitations in this study. (1) In spite of the usage of multiple Omicron-sensitive hACE2 transgenic rodent models in our study which revealed the attenuated pathogenicity of JN.1, it is unclear whether the JN.1 causes less clinically severe respiratory disease than previously circulating VOIs in humans. (2) Although the hamster model is the most widely used to evaluate the transmission of SARS-CoV-2, the household transmission of JN.1 was not known, especially when pre-immunity against multiple subvariants were taken into consideration. Clinical studies are needed to corroborate our findings in the transgenic rodent models.

## Materials and Methods

### Epidemiological

Monthly frequencies over one year of the indicated lineage in China, the United States, Europe and Singapore by August 2024 were downloaded from GISAID (https://www.epicov.org) on 30 August, 2024.

### Cells and viruses

Vero E6 cells and HeLa^hACE2+^ cells were cultured in Dulbecco’s modified Eagle’s medium (DMEM, Thermo) with 10% fetal bovine serum (Thermo) and 1% penicillin-streptomycin as described previously (15, 16). Vero E6^TMPRSS2+^ cells (from Japanese Collection of Research Bioresources Cell Bank) were cultured in DMEM with 10% fetal bovine serum, 1% penicillin-streptomycin and 1 mg/ml G418 (Sigma). The SARS-CoV-2 Omicron subvariants in this study include a JN.1 isolate (hCoV-19/Jilin/JSY-CC15/2024, GISAID Accession No. EPI_ISL_19231453) which contains two extra mutations C21254T (NSP16_A2596V), T21738C (Spike_F59S), a BA.2 isolate (hCoV-19/Jilin/JSY-CC5/2022, GISAID Accession No. EPI_ISL_18435548) and a EG.5.1 isolate (hCoV-19/Jilin/JSY-CC9/2023, GISAID Accession No. EPI_ISL_18908495). SARS-CoV-2 was passaged in Vero E6^TMPRSS2+^ cells, and the viral titers were measured using plaque forming unit (PFU) assays in Vero E6^TMPRSS2+^ cells.

### Western blot and viral load

The qRT-PCR assays were also used to determine the relative viral loads of SARS-CoV-2, as previously described (28). Tissues were lysed in RIPA buffer (Beyotime) supplemented with protease inhibitor cocktail (MCE) by homogenization in TissueLyser II (Qiagen). Tissue proteins were separated on precast SDS-PAGE (Epizyme). Western Blot was performed by using anti-hACE2 (Abcam) and anti-β-actin antibodies (CST).

### RNA sequencing

RNA sequencing was performed as previously described (29). Briefly, cellular RNA was extracted with an RNAsimple Extraction kit (Tiangen). After the RNA concentration and integrity were checked using a Qubit 2.0 (Thermo) and an Agilent 2100 Bioanalyzer (Agilent), sequencing libraries were generated using the NEBNext Ultra II directional RNA Library Prep Kit for Illumina (NEB) following the manufacturer’s recommendations. The library insert size was assessed on an Agilent 2100 system. The clustering of the index-coded samples was performed on a cBot Cluster Generation System. After cluster generation, the library was sequenced on the Illumina NovaSeq 6000 platform, and 150 bp paired-end reads were generated. Reads were mapped to the reference mouse genome (GRCm39). Differences in the expression of RNAs were evaluated on the basis of FPKM values with the software RSEM. Differential expression was determined by comparing virus-infected replicates (n=4) to mock-treated replicates (n=3) according to the criteria of an absolute log2FC > 1 and a false discovery rate (FDR)-adjusted P-value < 0.05.

### Pathogenicity

To reveal pathogenicity in mice, nine 7-week-old K18-hACE2 (K18-hACE2-2A-CreERT2) mice (Cyagen Biosciences) per group, including five animals in the body weight subgroup and four animals in the euthanized subgroup for tissue sample collection, were intranasally inoculated with the indicated virus at a dose of 1000 PFU (in 50 μl).

To reveal pathogenicity in hamsters, 52-week-old H11-K18-hACE2 hamsters (State Key Laboratory of Reproductive Medicine and Offspring Health, China) were intranasally inoculated with the indicated virus at a dose of 1000 PFU (in 100 μl). Each infected group included four animals in the body weight subgroup and five animals in the euthanized subgroup for tissue sample collection.

### Histopathology

The lungs of the animals were fixed in 4% paraformaldehyde in phosphate-buffered saline (PBS) and processed for paraffin embedding. The paraffin blocks were sliced into 3-µm thick sections and mounted on silane-coated glass slides, which were subsequently subjected to H&E staining for histopathological examination. Pathological features, including (i) bronchiolitis or exfoliation of epithelial cells in terminal bronchioles; (ii) hemorrhage; (iii) alveolar damage, including alveolar thickening or disappearance; and (iv) inflammatory infiltration, were scored using a four-tiered system as 0 (negative), 1 (weak), 2 (moderate), or 3 (severe). Tissue sections were processed for IHC with an anti-nucleocapsid antibody (Genetex) and a horseradish peroxidase-conjugated secondary antibody and finally stained with a DAB substrate kit (ZSbio). Panoramic images of the digital slides were taken using an WS-10 slide scan system (Wisleap).

### Airborne transmission

In the studies of airborne transmission between hamsters, we used six-week-old male WT hamsters or 52-week-old H11-K18-hACE2 male hamsters, respectively. Donor hamsters were firstly intranasally inoculated with virus at a dose of 1000 PFU (in 100 μl). After 24 hours, each infected donor hamster was cohoused with one corresponding acceptor hamster in isolation devices. The propagation cages containing either a donor or an acceptor were separated by 3 cm to prevent direct contact. Tissue samples were collected two days after infection for the donors or two days after the initial cohousing for the exposed acceptors.

### Nasal washes and exhaled aerosols

To collect nasal washes in another independent experiment, five wild-type hamsters were challenged with the indicated virus at a dose of 1000 PFU (in 100 μl). After collecting the nasal washes from 1-3 DPI, we measured the viral loads and viral titers.

Exhaled aerosols were collected in our published study (1). Specifically, exhaled virus aerosols from each group containing four golden hamsters were collected via an Andersen six-stage sampler (TE-20-800, TISCH, Cleves, OH, United States). The sampling flow rate was 28.3 L/min, and the sampling times were 1 and 2 DPI. The Andersen six-stage sampler was graded according to aerodynamic particle size, specifically: ≥7.0 μm, 4.7–7.0 μm, 3.3–4.7 μm, 2.1–3.3 μm, 1.1–2.1 μm, and 0.65–1.1 μm. Before each sampling session, the samples were fully treated with 75% alcohol and a nucleic acid scavenger (R504-01, Vazyme, China) and thoroughly dried. Viral aerosols exhaled were collected in pre-sterilized gelatin filters (12602, Sartorius, Germany). Each gelatin filter membrane was equally divided into six portions, three of which were used for viral titer titration. The remaining three portions were used for viral load detection.

For all animals, intranasal inoculation and euthanasia were performed under isoflurane anesthesia.

### Statistical analysis

All the data are presented as the means ± SDs. Comparisons were performed using Student’s tests. Significance was defined as P<0.05 (*), P<0.01 (**) or P<0.001 (***) and was indicated either in the figures or mentioned separately in the text. The number of repeats is specified in individual panels using discrete points. All presented data are biological replicates.

### Ethics and biosecurity

All animal experiments were approved by the Animal Care and Use Committee of the Changchun Veterinary Research Institute (approval number: AMMS-11-2023-032). All experiments involving infectious SARS-CoV-2 were performed at the Animal Biosafety Level 3 Laboratories of the Changchun Veterinary Research Institute, Chinese Academy of Agricultural Sciences.

## Acknowledgments

This work is supported by the National Key Research and Development Program of China (2021YFC2302405 to X.W., 2023YFC0871100 to Y.G. and 2021YFF0702500 to J.L.).

## Author contributions

R.L., Q.J., C.Z., W.W., and J.X. conducted the experiments; H.Z., B.L., F.Y., X.X., J.L. and Y.G. provided the resources; X.W. designed the study; X.W., R.L. and Q.J. wrote the manuscript. Y.G., J.L., F.Y., and X.X. revised the first draft and approved the submitted version.

## Supplemental information

Figs. S1 to S5 are obtained from Supplemental Information 1.

